# Coenzyme A binding sites induce proximal acylation across protein families

**DOI:** 10.1101/2022.05.24.493335

**Authors:** Chris Carrico, Andrew Cruz, Marius Walter, Jesse Meyer, Cameron Wehrfritz, Samah Shah, Lei Wei, Birgit Schilling, Eric Verdin

## Abstract

Lysine Nε-acylations, such as acetylation or succinylation, are post-translational modifications that regulate protein function. In mitochondria, lysine acylation is predominantly non-enzymatic, and only a specific subset of the proteome is acylated. Coenzyme A (CoA) can act as an acyl group carrier via a thioester bond, but what controls the acylation of mitochondrial lysines remains poorly understood. Using published datasets, here we found that proteins with a CoA-binding site are more likely to be acetylated, succinylated, and glutarylated. Using computational modeling, we show that lysine residues near the CoA-binding pocket are highly acylated compared to those farther away. We hypothesized that acyl-CoA binding enhances acylation of nearby lysine residues. To test this hypothesis, we co-incubated enoyl-CoA hydratase short chain 1 (ECHS1), a CoA-binding mitochondrial protein, with succinyl-CoA and CoA. Using mass spectrometry, we found that succinyl-CoA induced widespread lysine succinylation and that CoA competitively inhibited ECHS1 succinylation. CoA-induced inhibition at a particular lysine site correlated inversely with the distance between that lysine and the CoA-binding pocket. Our study indicated that CoA acts as a competitive inhibitor of ECHS1 succinylation by binding to the CoA-binding pocket. Together, this suggests that proximal acylation at CoA-binding sites is a primary mechanism for lysine acylation in the mitochondria.

## Introduction

Lysine acylations, such as acetylation, succinylation, or glutarylation, are post-translational modifications (PTM) (Rardin et al. 2013; Sadhukhan et al. 2016; Tan et al. 2014) that inhibit the actions of proteins across all kingdoms of life and all cellular compartments (Finkemeier et al. 2011; Weinert et al. 2013; Weinert et al. 2014). In eukaryotic cells, acylation of histones diminishes the electrostatic affinity between histones and DNA and is generally associated with an increase in gene expression (de Ruijter et al. 2003). Coenzyme A (CoA) is a metabolite required in a diverse range of metabolic processes, including the biosynthesis of fatty acids and ketone bodies, amino acid metabolism, fatty acid oxidation, and regulation of gene expression (Theodoulou et al. 2014; Carrico et al. 2018). In eukaryotes, CoA thioesters, such as acetyl-CoA, succinyl-CoA, and glutaryl-CoA, act as the sole cellular acyl group donors and react with lysine residues via both 1) enzymatic transfer mediated by acetyltransferase enzymes, such as p300 (Verdin and Ott 2015; Narita et al. 2019) and, 2) non-enzymatic mechanisms facilitated by high local concentrations of acyl-CoA species and high pH (Wagner and Payne 2013; Weinert et al. 2013). In the cytosol and nucleus, lysine acylation is primarily driven by acyltransferase enzymes, such as p300, and patterns of differential acylation have been attributed to the specificity and distribution of these enzymes. In the mitochondrial matrix, however, no universal acyltransferase enzyme has been identified, and mitochondrial acylation is thought to be mostly non-enzymatic (Weinert et al. 2017; Weinert et al. 2015). However, the distribution of acylated lysines in the mitochondria is not stochastic, with orders of magnitude differences in acylation found between sites (Weinert et al. 2015). Why some mitochondrial lysine residues are more susceptible to acylations than others remains an important unanswered question.

Mitochondrial metabolism depends on multiple acyl-CoA species that serve as key intermediates in critical pathways, such as the TCA cycle (acetyl-CoA, succinyl-CoA), fatty acid oxidation (acetyl-CoA, propanoyl-CoA, longer-chain acyl-CoAs), ketone body catabolism (3-hydroxymethylglutaryl-CoA, acetoacetyl-CoA), and amino acid catabolism (succinyl-CoA, glutaryl-CoA, HMG-CoA). These reactive acyl-CoA species serve as acyl-donor for the non-enzymatic acylations of mitochondrial proteins (Wagner et al. 2017). Regulatory roles have been identified for a limited subset of lysine acylations, which coexist with the majority of lysine acylation that is non-regulatory and low-stoichiometry (Weinert et al. 2015; Weinert et al. 2017). Importantly, many mitochondrial protein lysine residues are not acylated, and subsequent measurements of acetylation stoichiometry in mouse liver show an extremely wide range of acetylation (Weinert et al. 2015). The goal of the present study is to investigate the molecular mechanisms that control the acylation of specific lysine residues in the mitochondria.

While no universal mitochondrial acyltransferases have not been identified, the deacylases that remove acyl groups from lysine residues are well characterized. In the mitochondria, human sirtuins SIRT3, SIRT4, and SIRT5 have been identified as the main lysine deacylases. Sirtuins are a family of conserved protein deacylases that use NAD as co-substrate (Chang and Guarente 2014; Verdin 2015). SIRT3 regulates protein acetylation (Schwer et al. 2002; Onyango et al. 2002), and SIRT5 regulates acidic acyl modifications, such as succinylation, malonylation, and glutarylation (Du et al. 2011; Zhang et al. 2011; Peng et al. 2011; Tan et al. 2014). Less is known about SIRT4 in terms of its enzymatic activity; however, it acts on branched-chain acyl-lysine residues (Anderson et al. 2017). Mass spectrometry studies have mapped the landscape of multiple acylation marks in multiple species, tissues, and subcellular compartments. Proteomic surveys of mitochondrial protein acylation using knockout mouse strains against the mitochondrial sirtuins SIRT3 and SIRT5 identified the sites of protein acetylation, succinylation, and glutarylation. As mitochondrial sirtuin knockout mice are phenotypically normal, they have been used to generate a high-quality proteomic map of acylated lysines in cell lines and tissues (Huynh et al. 2019; Rardin et al. 2013a; Sadhukhan et al. 2016). Despite the important role played by histone acetylation for gene regulation in the nucleus, acylomic studies revealed that the majority of cellular acylation occurs within the mitochondria (Kim et al. 2006; Zhao et al. 2010; Park et al. 2013), most likely through non-enzymatic transfer of acyl groups from CoA thioesters. CoA concentrations can reach from 2.2 to over 5 mM in the mitochondria, and cytosolic concentrations are estimated at 0.02–0.14 mM (Leonardi et al. 2005).

Interestingly, metabolic pathways (e.g., TCA cycle, fatty acid oxidation, ketone body catabolism, amino acid catabolism, and ketone body synthesis) containing enzymes that directly interact with reactive CoA species are enriched in acylations (Nishida et al. 2015; Rardin et al. 2013; Weinert et al. 2013). Often, experimentally validated sites occur on enzymes binding one or more acyl-CoA species, with inhibitory acylation found in or near the CoA binding site (Sadhukhan et al. 2016; Rardin et al. 2013; Wang et al. 2019). Whether CoA binding and enzyme lysine acylation are causally linked has, however, not been reported. Additionally, a recent study showed a modest increase in acetylation near nucleotide-binding sites and suggested that enzymes with ADP-binding Rossmann fold motifs may bind acyl-CoAs (James et al. 2020). The authors also observed an increase in lysine malonylation by malonyl-CoA near the nucleotide-binding site of glutamate dehydrogenase (James et al. 2020). These studies suggest that interactions with CoA species specifically enhance the acylation of CoA-binding proteins (CoABPs).

Here, we investigated the factors that control lysine acylation in the mitochondria. Using computational analyses of published acylomic datasets, we showed that CoABPs are approximately three times more likely to be acylated than non-CoABPs in the mitochondria. In addition, we modeled possible CoA structural conformations and found that, in CoABPs, lysine residues near the CoA-binding pocket are more likely to be acylated than those far away from the CoA-binding pocket. We hypothesized that acyl-CoA binding could enhance the acylation of nearby lysine residues (**Figure 1**) and tested this hypothesis on enoyl-CoA hydratase short chain 1 (ECHS1), a mitochondrial protein with a CoA-binding pocket. Incubation with succinyl-CoA induced widespread succinylation of most ECHS1 lysine residues. Importantly, succinylation was competitively inhibited by co-incubation with CoA and CoA inhibition at a particular lysine site was inversely correlated with the distance between this lysine residue and the CoA-binding pocket. This finding suggests that proximal acylation at protein CoA-binding sites is a primary mechanism for lysine acylation in ECHS1 and likely more generally in the mitochondria.

**Figure 1.**
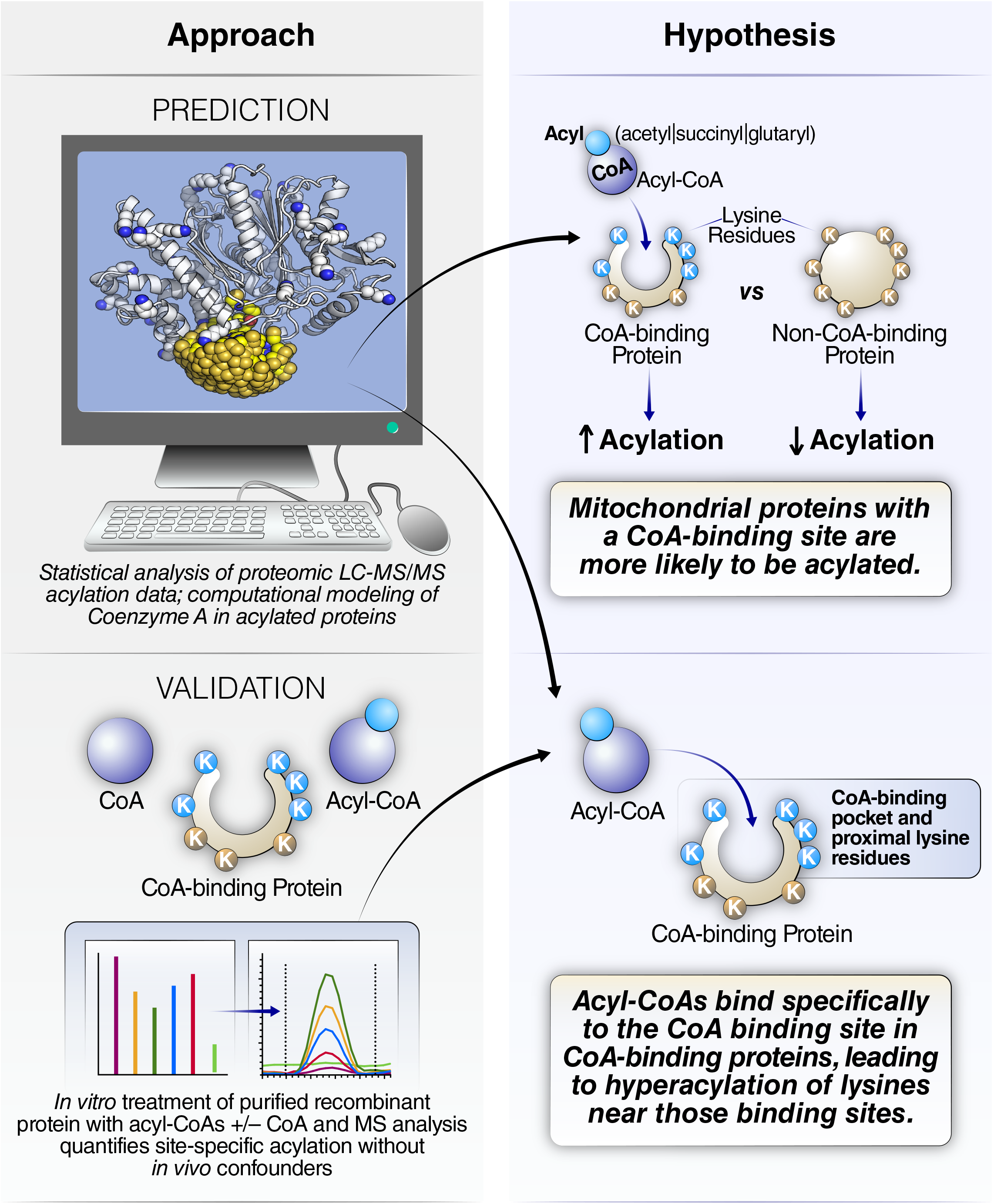
Computational modeling and mass spectrometry analysis revealed the mechanism of lysine acylation on CoA-binding protein within the mitochondria. (a) Schematic diagram of the experimental approaches in this study (left side) and the hypotheses tested (right side). Statistical analyses of proteome-wide MS acylation datasets using database annotation data, and subsequent structural modeling of acylated proteins (top left), generated a proposed mechanism for CoA-binding-protein hyperacylation (right). This mechanism was validated by *in-vitro* MS analysis of a reported acylated protein (bottom left).

## Results

### CoABP lysines are enriched in mitochondrial acylomic datasets

CoA thioesters are the major acyl group donor in mammalian cells (Carrico et al. 2018; Baldensperger and Glomb 2021) (**Figure 2a**). We hypothesized that CoABPs should be enriched in acylation marks, especially in the mitochondria where CoA thioesters are present in high concentrations (**Figure 1**). To determine if CoA binding induces lysine acylation, we calculated the fraction of acylated lysines identified in CoABPs among all acylated lysines. A list of CoABPs was generated from Uniprot annotation data **(Supplementary Table 1**). In the annotated mouse proteome, 2.7% of lysine residues belong to CoABPs (14,759 out of 547,206). By contrast, in a whole mouse liver acetylation dataset (Svinkina et al. 2015), 26.5% of acylated residues were from mouse CoABPs (512 out of 1,934) (**Figure 2b**). Calculating a relative odds ratio (OR) showed that lysines on CoABPs were 12.99 times more likely to be acetylated than lysines on non-CoABPs (OR=12.99, p = 3.68E-297).

**Figure 2.**
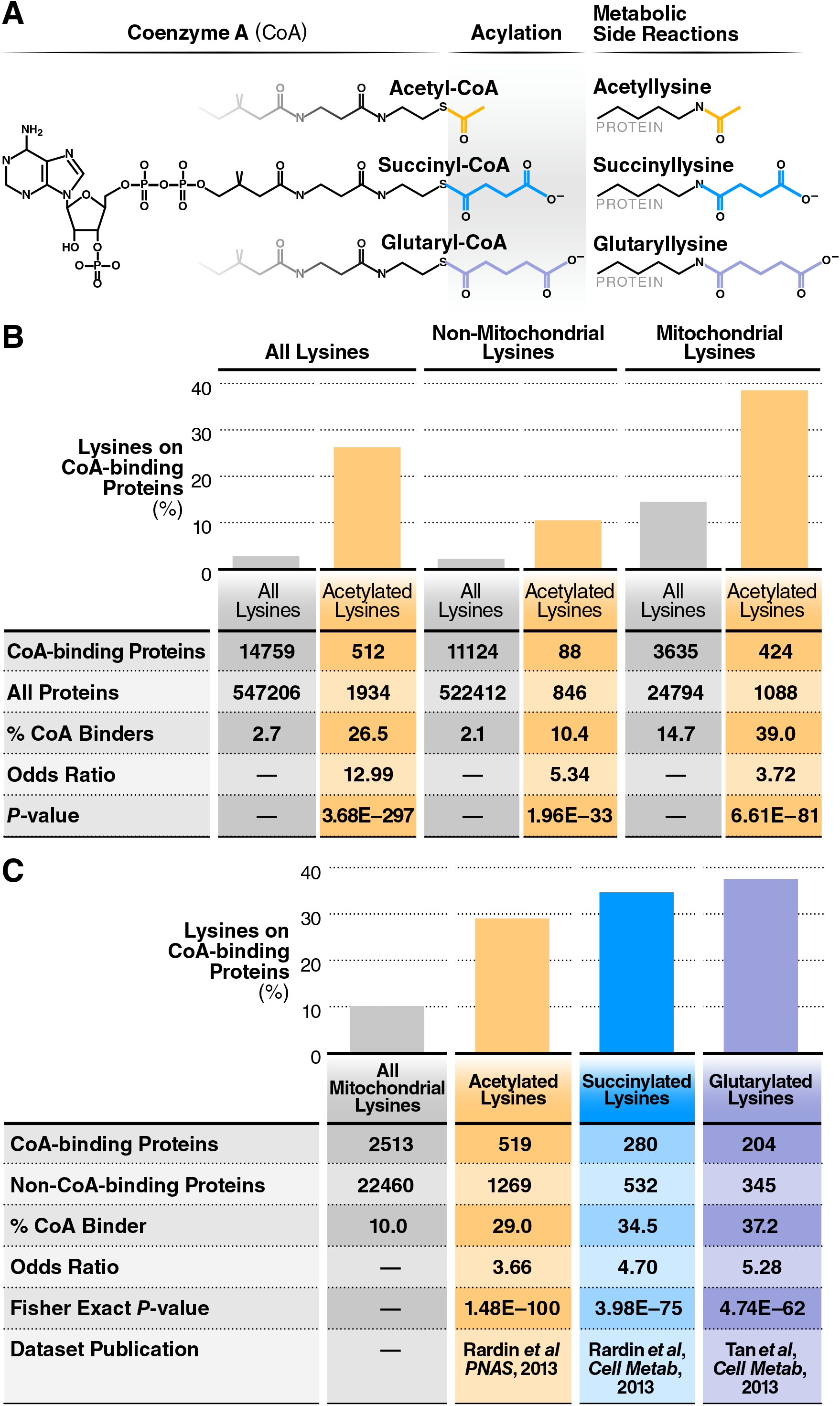
Lysine acylation is significantly over-represented on CoA-binding proteins. (a) Diagram of acetylation, succinylation, glutarylation modifications to protein lysine residues. (b) Table showing the enrichment of acetylated lysines on CoA-binding proteins analyzed in a whole-cell mouse liver acetylome dataset. This enrichment is also present when the dataset was separated into non-mitochondrial and mitochondria lysines. Significance was calculated using Fisher’s exact test. (c) Table showing the enrichment of acetylated, succinylated, and glutarylated lysines on CoA-binding proteins, using mitochondrial mouse liver datasets. For (b) and (c), statistical significances were calculated using Fisher’s exact test.

Since the majority of cellular acylation occurs in the mitochondria (Kim et al. 2006; Park et al. 2013), we compared lysines coming from mitochondrial and non-mitochondrial proteins (**Figure 2b**). Lysines annotated in CoABPs represented 14.7% and 2.1% of all lysines in mitochondrial and non-mitochondrial proteins, respectively. In the whole mouse liver dataset referenced above, 39.0% (424 of 1088) of mitochondrial acetylated lysines and 10.4% (88 of 846) of non-mitochondrial lysines belonged to CoABPs (**Figure 2b**). Lysines on CoABP were 3.72 times more likely to be acetylated than non-CoABP lysines in the mitochondria (OR=3.72, p = 6.61E-81), and 5.34 times more likely outside of the mitochondria (OR=5.34, p = 1.96E-33) (**Figure 2b**). We observed a similar enrichment in a whole-cell acetylation dataset in human Hela cells, albeit at a lower level (Hansen et al. 2019). In Hela cells, lysines on CoABPs were 1.92 more likely to be acetylated than non-CoABP lysines (OR=1.92, p = 3.61E-61, **Supplementary Figure S1**).

Most cellular acylation is found in the mitochondria, and we focused the rest of our investigation on mitochondrial proteins, using acylomic datasets from mitochondria extracts. Using the same approach, we determined if other types of acylation were enriched in mitochondrial CoABPs from three separate acylomic datasets. Mitochondrial lysine acylations, including acetylation (Rardin et al. 2013a), succinylation (Rardin et al. 2013b), and glutarylation (Tan et al. 2014) (**Figure 2a**), have been mapped in knockout strain mouse tissue samples lacking the corresponding sirtuin deacylases that remove those modifications. Among mitochondrial proteins, lysine residues in CoABPs represented 10.1% of all lysines (2,513 out of 24,973). By contrast, acetylated lysines in CoABPs represented 29.0% of all acetylated lysines (519 out of 1,788, **Figure 2c**). Similarly, in mouse liver mitochondrial samples, succinylated and glutarylated lysines in CoABPs represented, 34.5% (280 out of 812) and 37.2% (204 out of 549) of all succinylated or glutarylated proteins, respectively. (**Figure 2c**). In these mitochondrial datasets, lysines on CoABPs were 3.66, 4.70, and 5.28 times more likely to be acetylated, succinylated, or glutarylated than lysines on non-CoABPs (OR=3.66, p = 1.48E-100; OR=4.70, p = 3.98E-75; OR=5.28, p = 4.74E-62, respectively). Thus, in the mitochondria, lysine residues in CoABPs are significantly more likely to be acylated than those in non-CoABPs.

### Simple CoA conformational model shows significant acylation enrichment on CoA-accessible lysine residues

Next, we hypothesized that CoA binding itself leads to lysine acylation and, as a consequence, lysines near a CoA-binding site are acylated with a higher probability than those distal to CoA-binding sites (**Figure 1**). To test this hypothesis, we used crystal structures of CoA-binding proteins and computational modeling of CoA binding at structurally defined sites to determine if proximal lysines were enriched in acylation.

Within the CoA molecule, the acyl group is conjugated to a thiol group linked to a core phospho-ADP moiety by a ∼15Å prosthetic group with 11 rotationally variable sites (**Figure 3a**). The terminal acyl group of this large, flexible molecule can assume multiple positions relative to the core phospho-ADP moiety, and calculating the physical distance between a lysine residue and the acyl group of an acyl-CoA thioester bound to a protein is not straight-forward. Crystal structures are available for CoA bound to many protein active sites, and we leveraged this information to compute an ensemble of ∼2000 models of sterically valid CoA conformations (**Figure 3a, supplemental code text**). These models were selected to achieve the largest possible spread of terminal CoA sulfur positions. AMP is the largest conformationally rigid subsection of the CoA molecule, and so, we aligned this conformational ensemble to homology models of CoABP, using the AMP moiety as a target and the subset of our CoA ensemble conformations that sterically fitted against each protein model (**Figure 3a**). Each protein’s ensemble of sterically valid CoA conformations describes a set of thiol sulfur positions that, in turn, define a set of distances to each potentially acylated lysine residue’s amine group. For each CoABP analyzed, the minimal distance between each lysine residue from the fitted CoA conformational ensemble was calculated. (**Figure 3a**).

**Figure 3.**
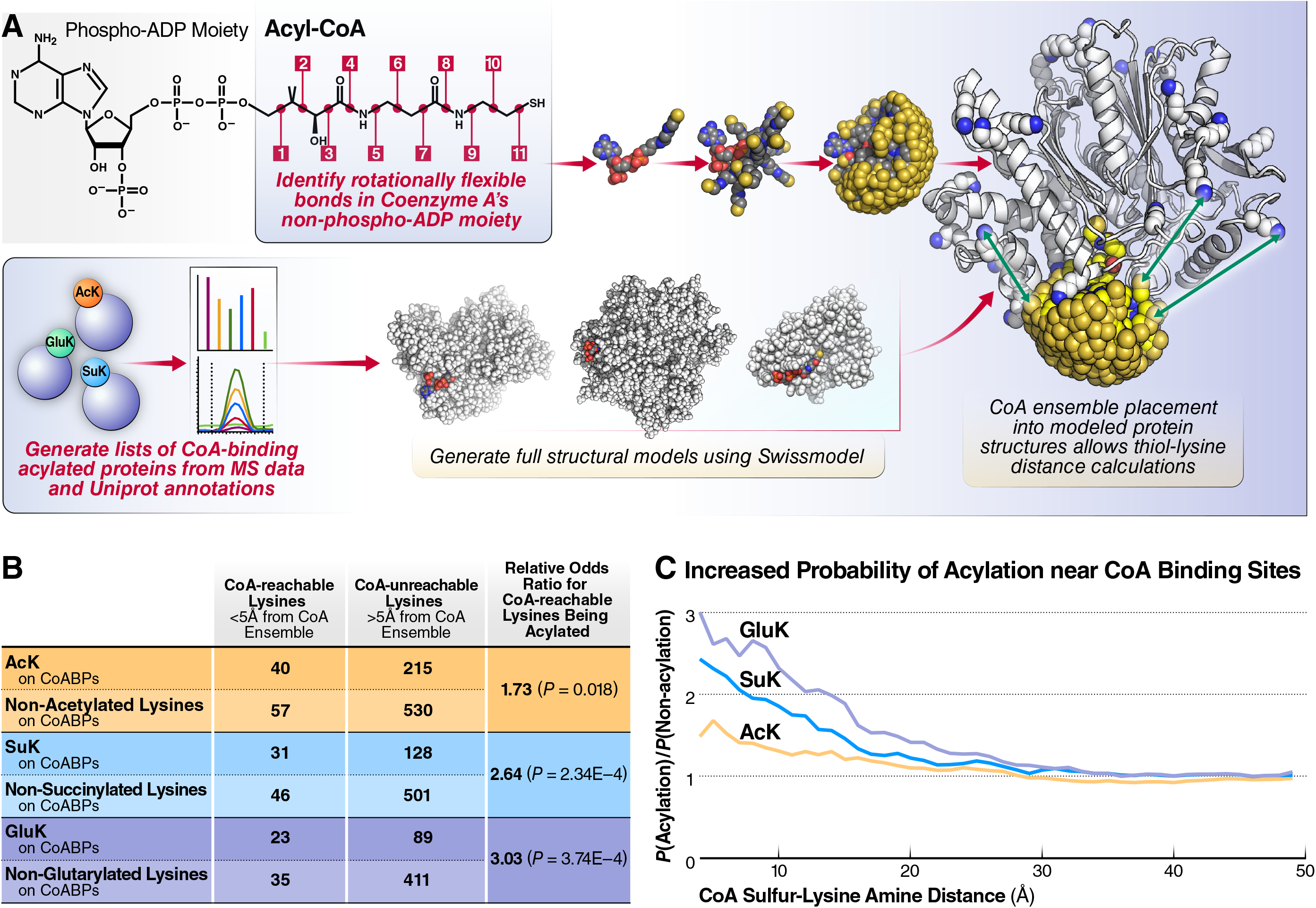
Structural modeling of CoA conformation reveals proximal hyperacylation near CoA-binding sites. (a) Schematic diagram of the computational approach used to calculate the minimal distance between a lysine residue the CoA’s thiol group in a protein CoA-binding pocket. (top) A conformational ensemble of physically plausible CoA conformation was made by combining experimentally observed bond rotational angles from CoA crystal structures after the phospho-ADP moiety. (bottom) Full structural models of CoA-binding proteins from acylomic datasets were generated using Swissmodel. (right) The CoA ensemble was docked into each protein model to generate a set of sterically accessible CoA thiol locations and used to score each modeled lysine residue’s amine-thiol distance. (b) Table showing the numbers of non-acylated and acylated lysine residues found on CoA-binding proteins. Relative odds ratio for CoA-reachable lysines (<5Å from the nearest CoA ensemble sulfur) being acylated vs. more distal lysine residues were calculated for each of acetylation, succinylation, and glutarylation. The significance of the results was calculated using Fisher’s exact test. (c) Relative probability of lysine acylation on CoA binding protein lysines as a function of a lysine’s distance from the CoA ensemble. Data shown for acetylation, succinylation, and glutarylation.

We used a distance of 5Å between the lysine amine and the CoA sulfur as the cutoff for close binding site proximity and again modeled acylation observations as an odds ratio. That distance was empirically chosen to compensate for limited CoA sulfur position sampling density and the rejection of CoA sulfur positions with Van der Waals overlap to lysine amine atoms (Smith and Denu 2009). The OR values for CoA-reachable lysines being acetylated, succinylated, and glutarylated were 1.73, 2.62, and 3.03, respectively, and those increased acylation odds were all significant (**Figure 3b**). By plotting the cumulative fraction of all modeled CoABP lysines within a given distance of the CoA ensemble, we observed that lysines close to the CoA binding site were 2–3 times more likely to be acetylated, succinylated, and glutarylated than lysines farther away (**Figure 3c**). As the distance increased, the probability of acylation decreased down to background levels (**Figure 3c**). These results validated our hypothesis and indicated that lysine acylation was significantly increased near CoA-binding sites.

### Acylation enrichment near the CoA-binding site is specific to CoA

We next wanted to verify that the increase in acylation near CoA-binding sites was specific to CoA. CoA shares a core ADP moiety with related molecules (e.g., ADP, ATP, NAD, and NADP) (**Figure 4a**). However, the final step of CoA synthesis adds a phosphate to its ribose group in a position unique to CoA, potentially sterically excluding CoA from the binding sites used by these related molecules. The 3′-phosphate CoA molecule is typically bound by a highly positively charged binding site, requiring either lysine or arginine residues in protein sequences and leading to a statistical enrichment of lysines near CoA binding sites (Bharathi et al. 2013; Choosangtong et al. 2015). Since the related molecules have sizes and polyvalent negative charges similar to CoA, their binding sites also depend on positively charged amino acids. Given the increase in acylation near CoA binding sites (**figure 3b** and **3c**), local charge effects may drive nonspecific CoA binding and subsequent acylation. To rule out this possibility, we compared acylation near NAD/NADP binding sites with CoA-binding sites to determine if the local charge effect is sufficient to drive local acylation (**Figure 4a**). We modeled CoA steric accessibility for NAD[P] sites, using the same protocol as CoA and the shared ADP moiety as the base for determining binding site proximity for lysines on NAD[P]-binding proteins (**Figure 4b,4c**). Again using 5Å as a cutoff for close binding, we observed no higher probability of glutarylation in CoA-reachable lysines in NAD[P]-binding proteins since no glutarylated lysine was actually ‘CoA-reachable’ (**Figure 4b**). Unlike CoABPs, where acylation probabilities significantly increased with the proximity of the lysine to a CoA-binding site, no significant changes in acylation were observed on lysines close to NAD- and NADP-binding sites (**Figures 4b** and **4c**). Thus, acylation enrichment near CoA-binding sites is specific to CoA-binding sites and cannot be explained by a more generalizable charge effect.

**Figure 4.**
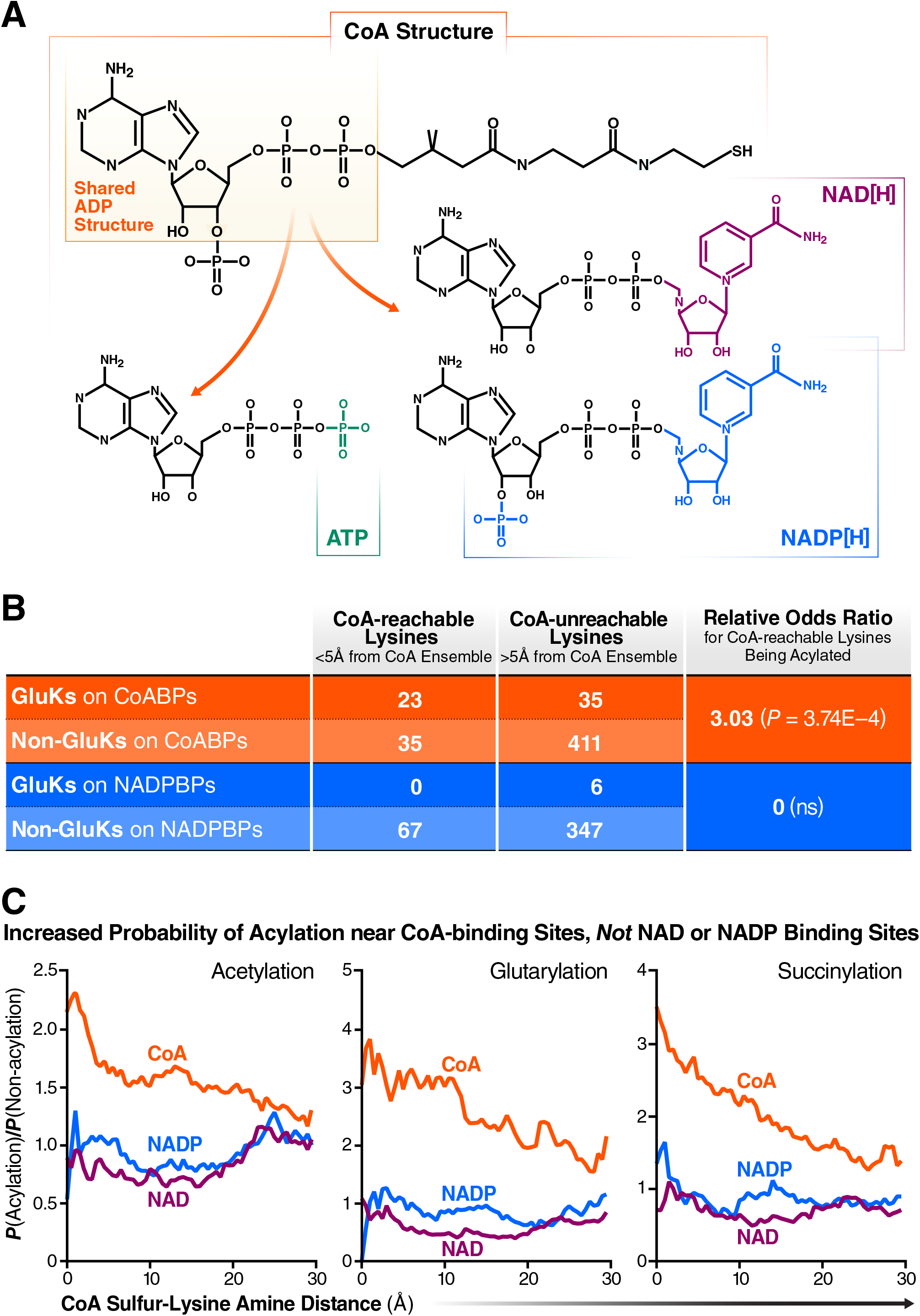
Proximal hyperacylation is specific to CoA-binding sites. (a) Illustration of the structural similarities between CoA and ADP, ATP, NAD, and NADP. CoA shares a core ADP moiety with ATP, NAD, and NADP, but its added 3′ phosphoryl group may prevent it from fitting in binding sites of those species. (b) Table comparing the relative odds ratios of CoA-reachable lysines being glutarylated by CoA-binding protein and NAD[P]-binding proteins. (c) Relative probability of lysine acylation as a function of a lysine’s distance from the CoA ensemble built into [acyl-]CoA, NAD, and NADP binding sites, showing a lack of acylation enrichment near NAD and NADP binding sites.

### Free CoA inhibits acylation of CoA-binding proteins

A likely mechanism that would explain our observations is that acyl-CoAs occupy CoA binding sites and transfer their acyl group to nearby lysine residues, either directly or using nearby cysteine residues as intermediates (James et al. 2018). In agreement with this model, incubation of CoABPs with acetyl-CoA or succinyl-CoA yields detectable hyperacetylation after a period of hours (Wagner and Payne 2013). If this hypothesis is correct, we can expect that 1) lysine residues near the CoA binding site should become acylated at a faster rate, and 2) acylation should be competitively inhibited by an excess of non-acylated CoA (**Figure 5a**).

**Figure 5.**
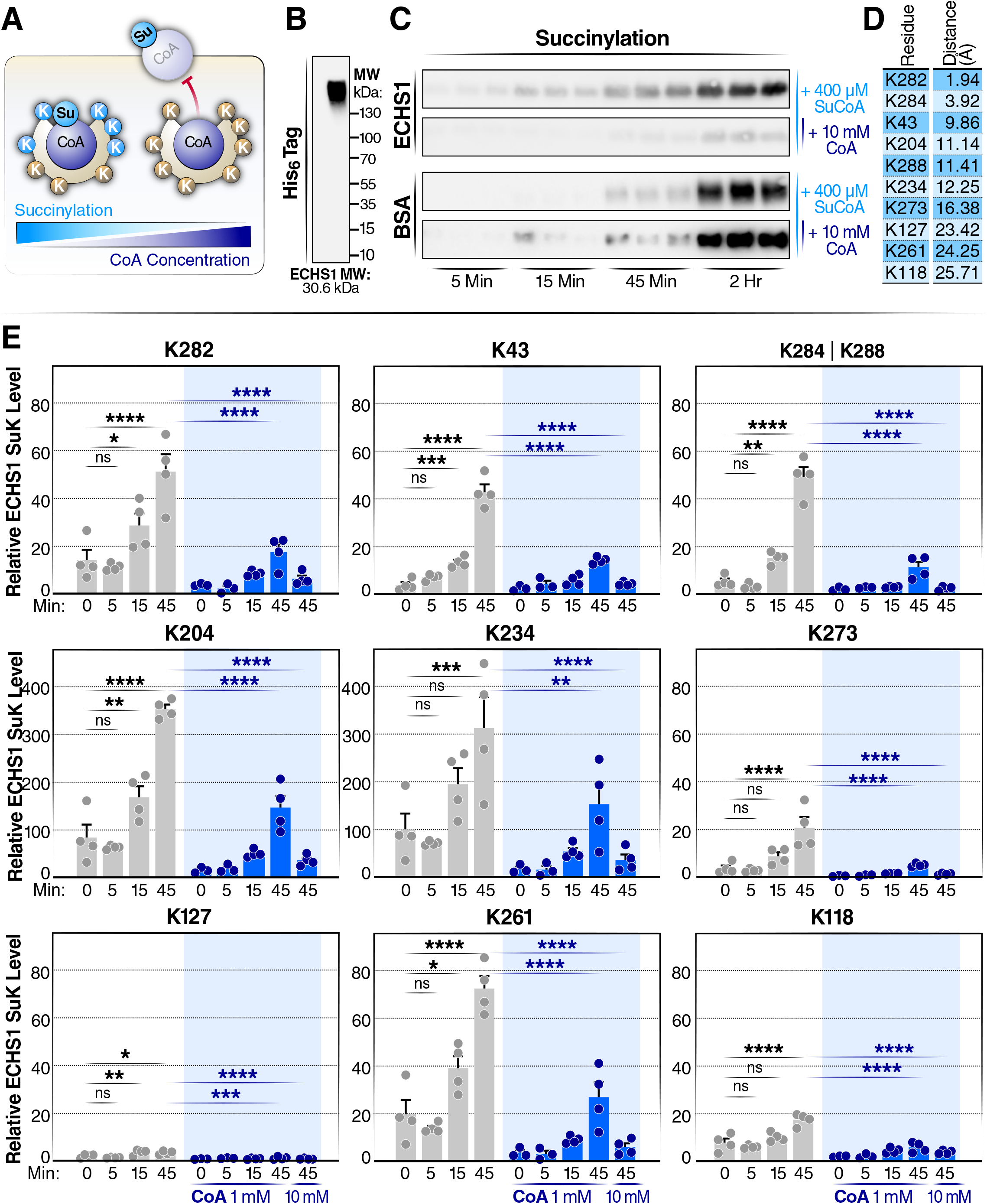
*In-vitro* succinylation of ECHS1 with succinyl-CoA is competitively inhibited by CoA. (a) Schematic diagram of the succinylation of ECHS1 by succinyl-CoA (Su-CoA) and inhibition by CoA. (b) Western blot of human recombinant ECHS1 using native gel electrophoresis. (c) Western blot detection of succinyl lysine (SuK) levels in ECHS1 and BSA co-incubated with Su-CoA in the presence and absence of CoA for varying times (5, 15, and 45 min, and 2 h). (d) Distance of individual lysine residues to CoA on ECHS1. (e) Mass spectrometry time-course experiment measuring the change in succinylation levels at individual lysines on ECHS1. ECHS1 was co-incubated with 400 µM Su-CoA for 0, 5, 15 or 45 min and with 0, 1 or 10 mM CoA. The abundance of SuK levels for each residue was normalized to the ECHS1 negative control (not treated with CoA or Su-CoA), (N = 4 per treatment). ^*^;P<0.05, ^**^P<0.01, ^***^P<0.001 and ^****^P<0.0001; ns, not significant. A one-way ANOVA was performed. In the conditions not treated with CoA, each time point was compared to the 0 min group. In the 45-min conditions, both CoA-treated groups (1 and 10 mM) were compared to the group not treated with CoA.

We sought to test this hypothesis using recombinant enoyl-CoA hydratase, short chain 1 (ECHS1), purified from *E. coli*. ECHS1 is a 30.6-kDa mitochondrial CoABP. ECHS1 forms homohexamers, and native-gel electrophoresis confirmed that purified ECHS1 was present in a highly multimerized state (**Figure 5b**). Since endogenous acetylation often occurs on bacterial-expressed proteins (Weinert et al. 2013), we used succinyl-CoA as our model acyl-CoA for most of the experiments. ECHS1 was incubated with succinyl-CoA (400 μM), and we measured how increasing incubation time (5, 15, 45, and 120 mins) affected the succinylation of ECHS1.

Bovine serum albumin (BSA) was used as a non-CoABP control. Western blotting results showed that succinylation levels induced by succinyl-CoA incubation were increased in a time-dependent manner in both proteins (**Figure 5c**). Importantly, co-incubation with a 25-molar excess of CoA (10 mM) strongly inhibited succinylation of ECHS1, but not of BSA. This suggests that succinylation by succinyl-CoA depends partially on its interaction with the ECHS1 CoA-binding pocket and that CoA inhibition is specific to CoABPs (**Figure 5a**).

To characterize the differentially succinylated lysine sites in detail, ECHS1 succinylation was analyzed by liquid chromatography, data-dependent acquisition tandem mass spectrometry (LC-DDAMS/MS) after protein gel-purification and in-gel trypsin digestion. We observed that 10 out of 24 lysine residues within ECHS1 (**Figure 5d**) were succinylated by incubating with succinyl-CoA. Of note, K284 and K288 within the same disuccinylated peptide cannot be distinguished. In line with the western blot results, the levels of succinylation induced by succinyl-CoA incubation were increased on each of these lysine residues (K282, K284/K288, K43, K204, K234, K273, K127, K261, K118) in a time-dependent manner at 0, 5, 15, and 45 min (**Figure 5e**). Moreover, succinylation of ECHS1 was inhibited by CoA at these lysine residues. Thus, succinylation of individual lysine residues on ECHS1 was inhibited by CoA in a concentration-dependent manner.

To determine the sensitivity of individual lysine residues to CoA inhibition, we incubated recombinant ECHS1 with succinyl-CoA (400 μM) and increasing concentrations of CoA (0–10 mM). After protein digestion and LC-MS/MS analysis, we obtained robust MS data for seven lysine residues (i.e., K282, K284/K288, K43, K204, K234, and K118). Importantly, succinylation was competitively inhibited by the increasing concentration of CoA (**Figure 6a**), and the IC_50_ for each of these inhibition curves ranged from 331 to1924 μM. Next, we asked, for each succinylated lysine residue, whether CoA-induced inhibition was affected by its distance to the CoA-binding pocket. Lysine distances to the CoA binding pocket were calculated with the same method described above (**Figure 3a**), using the ECHS crystal structure (PDB ID: 2HW5) and calculating the set of sterically accessible conformations for CoA. We plotted lysine distances from the CoA binding pocket with the IC_50_ values and performed a linear regression analysis.

**Figure 6.**
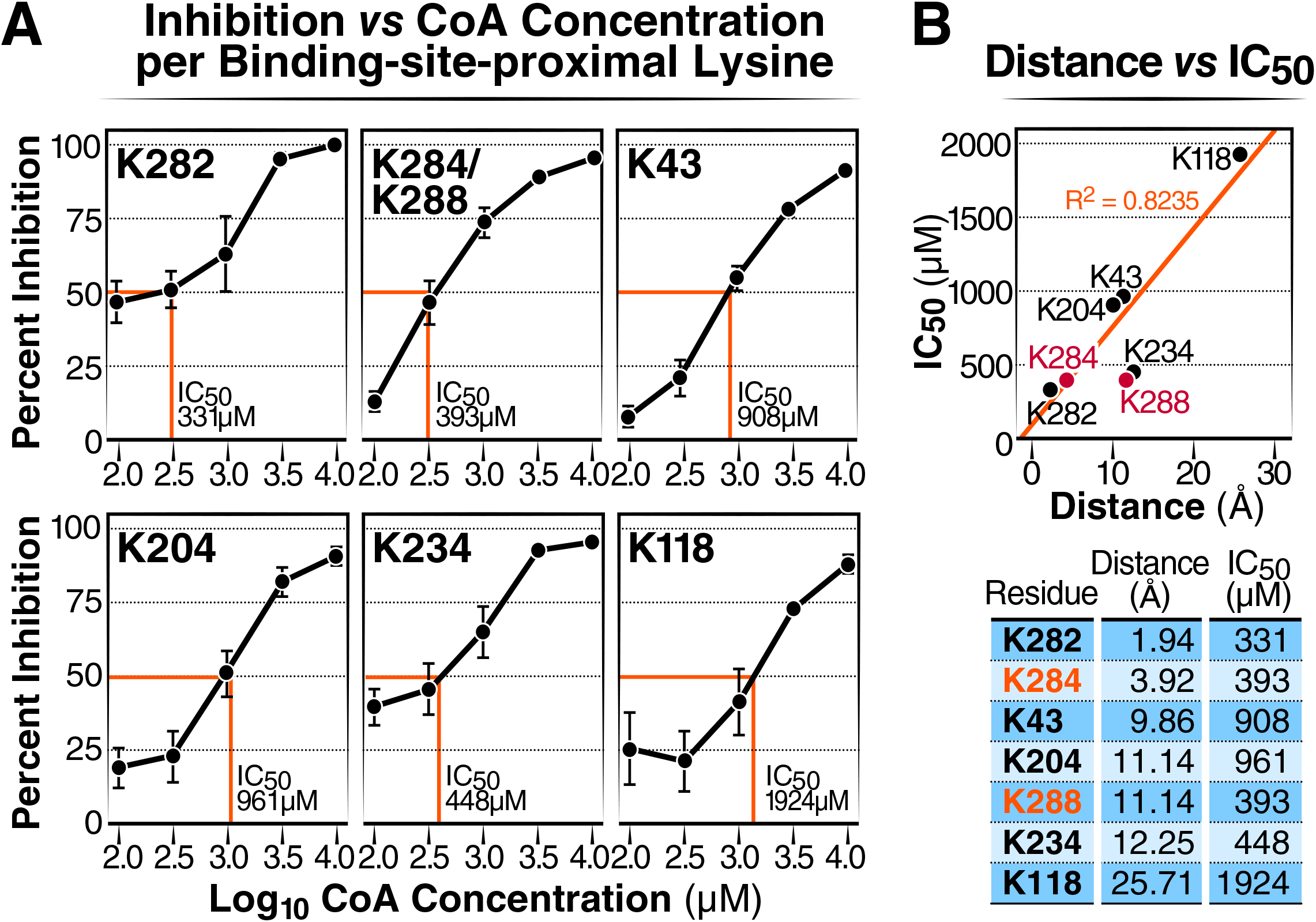
CoA inhibition at a particular lysine site is inversely correlated with the distance to the CoA-binding pocket. (a) Inhibition curve and per-residue IC_50_ for CoA inhibiting the succinylation of individual lysine residues on ECHS1. ECHS1 was incubated with succinyl-CoA and increasing concentrations of CoA. Dose-response curves were expressed as the log of CoA concentration vs SuK inhibition. Percent inhibition for each residue was calculated by measuring the percent change between succinylation levels when ECHS1 was co-incubated with succinyl-CoA in the presence and absence of CoA treatment for 45 min. N = 4 per treatment. (b) Plot showing the correlative relationship between lysine residue distances vs the IC_50_ (μM) (N= 5, R^2^= 0.8235). In the MS analysis, succinylation levels for K284 and K288 were combined as both residues were exclusively found on a shared peptide after trypsinization. As a result, these sites were not used to generate the R^2^ value or trendline and are included in the graph as red points.

We observed a positive relationship between lysine distance from the CoA-binding pocket and the IC_50_ of each residue (R^2^=0.8235) (**Figure 6b**). This showed that lysines closer to the CoA-binding pocket were more sensitive to CoA inhibition than lysine further away. Altogether, these experiments indicate that lysine succinylation near the ECHS1 CoA-binding pocket is mediated by acyl-CoA binding and is subject to competitive inhibition by CoA (**Figure 1**).

In addition to this succinylation experiment, we performed a similar acetylation experiment using recombinant ECHS1 protein incubated with acetyl-CoA and CoA. Since endogenous acetylation occurs on bacterially expressed proteins (Weinert et al. 2013), we only analyzed acetylation on three lysine sites near the CoA-binding pocket. Similarly, MS analysis results showed that acetylation at lysine residues near the CoA-binding pocket (K282, K284, and K204) in ECHS1 induced by acetyl-CoA incubation was inhibited by a 25-molar excess of CoA (**Figure 7**). These results suggest that CoA more generally inhibits acyl-CoA-induced lysine acylation near CoA-binding pockets for multiple distinct acylation species.

**Figure 7.**
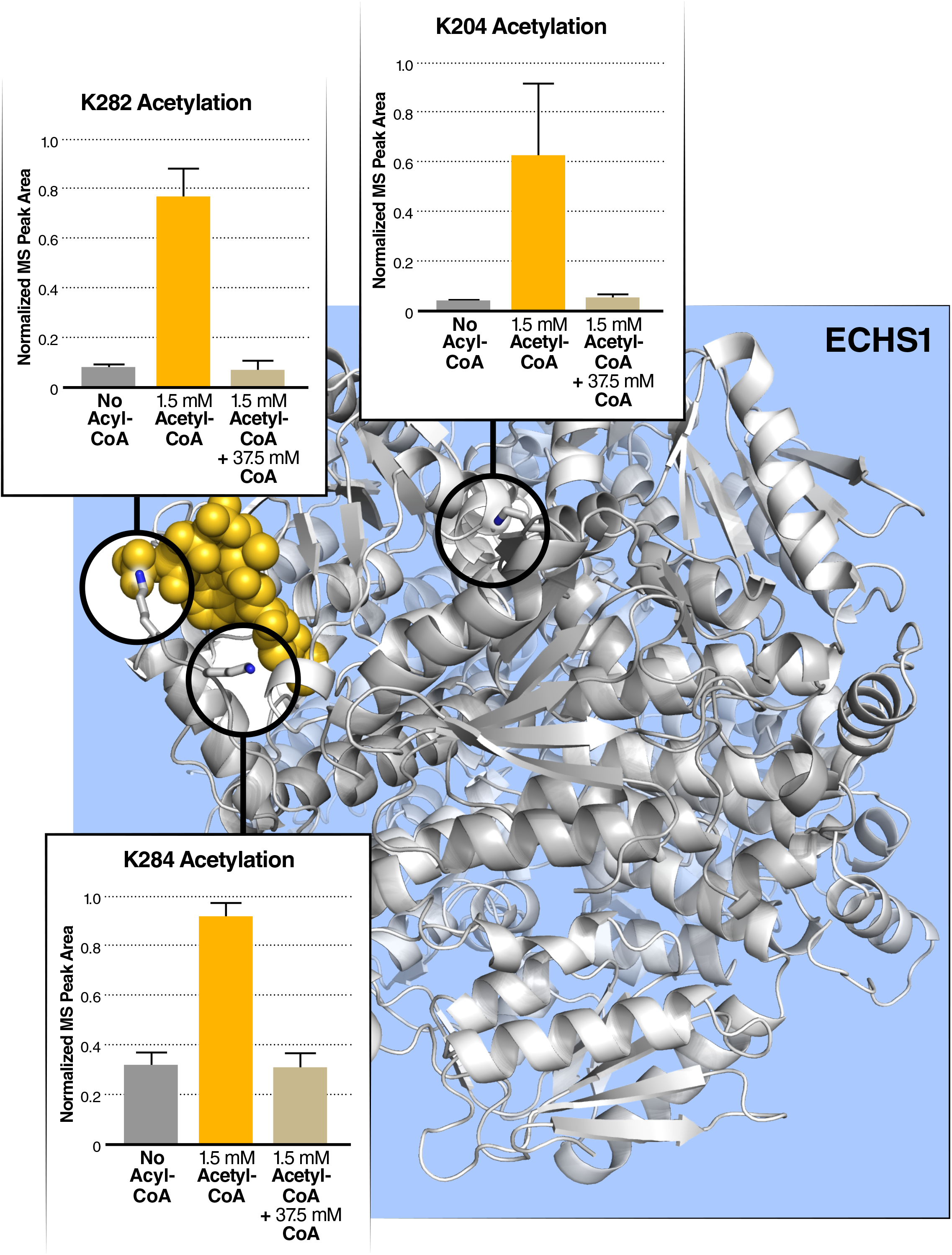
*In-vitro* acetylation of ECHS1 with acetyl-CoA is competitively inhibited by CoA at lysines proximal to the CoA-binding site. (a) Acetylation at three sites on bacterially produced recombinant ECHS1 increases *in vitro* after the addition of acetyl-CoA, and these sites are found near the CoA binding site. This acetylation via exogenous acetyl-CoA is inhibited by adding molar excess of CoA.

## Discussion

Using computational modeling validated by *in-vitro* experiments, we showed that acylation is enriched near the active site of CoABPs. In particular, our results indicate: (1) CoABPs are significantly more likely to be acylated, especially in the mitochondria, (2) lysines physically close to the CoA binding site are more likely to be acylated than those far away, (3) *in-vitro* acylation of CoABP depends on acyl-CoA binding, and (4) lysines near the CoA-binding site are more sensitive to inhibition than distal lysines. This study suggests proximal acylation at protein CoA-binding sites is a primary mechanism for lysine acylation in the mitochondria.

Protein modification reactions, such as phosphorylation, acylation, or glycosylation, are typically performed by dedicated transferase enzymes that stabilize reaction transition states and bring the modified protein and a small-molecule precursor together (Ohtsubo and Marth 2006; Singh et al. 2017; Wang and Lin 2021). In the case of binding-site-mediated proximal acylation, the reaction uses a preformed binding site for CoA (as with canonical enzymatic catalysis) but no specific transition site stabilization (as with non-enzymatic reactions). The CoA-mediated proximal acylation has features of both enzymatic and non-enzymatic reactions and could be qualified as a “semi-enzymatic” reaction mechanism, which enhances a class of related side reactions in a chemically selective and sterically constrained fashion.

CoA thioesters share both (1) a common binding moiety that can occupy a wide variety of binding sites throughout the proteome and (2) a highly reactive terminal bond that irreversibly transfers an acyl payload to a common nearby chemical group. These two factors generate a large number of detectable PTMs across the proteome and allow to perform the computational analyses described above. However, other reactive metabolites could exhibit the same behavior, albeit on fewer proteins. Other PTM-generating reactions, such as 1,3-bisphosphoglycerate (Moellering and Cravatt 2013), exhibit similar patterns of rate enhancement near binding sites for PTM precursor metabolites. Although the resulting set of PTMs on other reactive metabolites may not be sufficiently widespread for proteomic LC-MS/MS-based detection and analysis, more focused techniques may find inhibition of a wider set of metabolic enzymes as side effects of the reactions they catalyze. We believe that the combination of the computational and experimental approach used here uncovered new mechanisms in the field of acylation, and we anticipate that it also illuminates mechanisms for other PTMs induced by reactive metabolites or drugs.

In 2020, James et al. reported that enzymes incorporating the ADP-binding Rossman fold engage in autoacylation, similar to our observations (James et al. 2020). In particular, they showed that automalonylation by the NAD- and ATP-binding enzyme glutamate dehydrogenase (GDH) was (1) inhibited by a molar excess of bare CoA and (2) greater for lysine residues close to GDH’s nucleotide-binding sites. Using a linear distance function rather than a structurally modeled CoA ensemble, they also found that acetylation stoichiometry was, on average, modestly higher for lysine residues near nucleotide-binding sites on enzyme surfaces. Our study confirms and extends the validity of their finding. First, by considering multiple acylation subtypes in parallel, we see an unexpected increase in the enrichment for multivalent acidic modifications, such as glutarylation and succinylation, over acetylation. Second, while James et al. reported NAD-binding-site-driven automalonylation for GDH, we did not observe this effect at the proteome-wide level. They showed that NAD is an inferior inhibitor of autoacylation in mitochondrial protein extracts, compared with bare CoA, and that other ADP derivatives were intermediate in effect between NAD[P][H] and bare CoA. We suspect that, while some subset of NAD-binding sites binds CoA efficiently for autoacylation by acyl-CoA species, many others do not. GDH’s proclivity for inhibitory acylation may therefore be an outlier among NAD-binding enzymes. In contrast, every protein capable of binding CoA for its canonical function should exhibit some degree of autoacylation, though further study will be required to better understand differences in rates among enzymes as a consequence of their structure.

Finally, by computationally analyzing data from several hundred proteins in published datasets and validating our computational model using a single CoA-binding protein, we showed that acylation of mitochondrial CoA-binding protein occurs through acyl-CoA species binding to the CoA binding pocket. Further studies will be necessary to expand the validity of our findings, but with this publication, we hope that researchers interested in acylation can begin quantitatively analyzing the impact of protein biochemical and structural properties on the various acylations they accumulate. Ultimately, more detailed mechanistic models should also aid in identifying the metabolic conditions where these PTMs are most impactful on biology, as well.

## Materials and Methods

### Adjudication of CoA-binding proteins

Lists of observed peptides from published supplemental information from prior MS studies were used to generate UniProt IDs. A protein was considered to be CoA-binding if its enzymatic activity involved any CoA derivative, had any CoA derivative annotated as a ligand, or had an annotated CoA derivative binding or active site. Similar procedures were conducted for NAD- and NADP-binding proteins.

### Computational modeling of CoA ensembles in CoA-binding protein contexts

For each CoA- or NAD[P]-binding protein, homology models were generated using Swissmodel’s interactive model-building tools (Waterhouse et al. 2018). Proteins with homology models were subjected to structural alignment by PyMOL (The PyMOL Molecular Graphics System, Version 2.0 Schrödinger, LLC) against the set of experimentally determined crystal structures with CoA bound, using Python scripts to set up and compare alignments. Top pairs of acylated protein homology models and structurally similar CoA-bound proteins were used to generate CoA placements in the context of acylated homology models using only the CoA-local protein residues within 20Å of CoA phosphate atoms as a structural alignment target.

Once placed, the base phospho-AMP moiety from CoA was used as the alignment target to place a self-avoiding CoA conformational ensemble, from which each CoA confirmation was then excluded if it shared more than 1Å^3^ of collision volume with the protein model. This shared input CoA conformational ensemble was generated by applying backbone dihedral angles from existing CoA ligand structures from RCSB (Burley et al. 2021) to rotate an idealized CoA ligand structure. Individual conformations were generated by random sampling of experimentally measured backbones, and iteratively accepted into the final model if their terminal thiol was more than one sulfur atom Van der Waals radius away from any other model in the set until 2000 conformations had been selected.

For NAD[P] binding sites, the shared AMP core common to both CoA and NAD[P] was used to place the CoA conformational ensemble; modeling was otherwise performed just as for CoA-binding proteins.

Lysine residues on all homology models with built CoA ensembles were assigned a distance score from the center of each lysine’s terminal primary amine atom to the center of the nearest thiol sulfur atom among the model’s set of sterically valid CoA conformations.

For each lysine acylation subtype considered (acetylation, succinylation, and glutarylation), we evaluated lysine residues from all homology models where at least one modeled lysine residue was observed in the corresponding proteomic MS study. Individual lysine residues acylated vs. lysine residues not acylated among these acylated proteins were used in conjunction with the distance scores described above to calculate the relative enrichment of acylation within X Angstroms of the CoA ensemble, and Fisher’s Exact Test as implemented in Scipy.stats (Virtanen et al. 2020) was used to assess the statistical significance of enrichment.

Code developed for this study is available at https://github.com/ccarrico/CoABindingSiteAnalyses

### Protein incubations

Recombinant enoyl-CoA hydratase short chain 1 (ECHS1) (0.3 µg/µL) was purchased from Novus Biologicals and dissolved in 50 mM Tris, pH 8, 150 mM NaCl with varying concentrations of CoA (0–10 mM) at 37°C. The reaction was started by adding either succinyl-CoA or acetyl-CoA at 37°C. Reactions were halted by snap freezing in liquid nitrogen and stored at -80°C until processed.

### SDS-PAGE

Polyacrylamide gels (10–15%) were used for the separation of ECHS1 or BSA. The protein was transferred to nitrocellulose membranes using the BioRad blotting system. Membranes were blocked with 5% BSA in Tris-buffered saline (20 mM Tris, 150 mM NaCl, 0.1% Tween 20, pH 7.4) and probed with anti-succinyl lysine (PTM Biolabs), then probed with secondary antibody in milk in Tris-buffered saline, incubated with antibodies. Chemiluminescence intensities were detected using the ChemiDoc imaging system (BioRad, Hercules, CA).

### Mass spectrometry of ECHS1

For the MS experiments, 3 µg of ECHS1 was co-incubated with 400 µM succinyl-CoA for 0, 5, 15 or 45 min and with 0, 1, or 10 mM CoA (buffer: 50 mM Tris, pH 8.0, and 150 mM NaCl). Each condition contained four replicates.

#### Digestion

Each sample containing 3 µg of ECHS1 stock protein was incubated in 50 mM dithiothreitol (DTT) and Laemmli sample buffer sample buffer for 10 min at 70° C. The samples were run in precast 4–12% Bis-Tris stacked gels for 20 min. In-gel digestion was performed the following day. Gel bands were diced and collected in tubes and dehydrated with a dehydration buffer (25 mM ammonium bicarbonate in 50% acetonitrile and water). The gel samples were dried in the speed vac, reduced with 10 mM DTT and incubated for 1 h at 56° C with agitation, and then alkylated with 55 mM iodoacetamide and incubated for 45 min at room temperature in the dark. The diced gels were washed with 25 mM ammonium bicarbonate in water and then dehydrated once again with the dehydration buffer. The samples were dried in the speed vac once again after which the proteins were incubated in 250 ng of trypsin for 30 min at 4° C and digested overnight at 37° C with agitation. The following morning, the digests were subjected to water and then 50% acetonitrile and 5% formic acid in water. After each addition of solution, the aqueous digest from each sample was collected into a new tube. These pooled peptide extractions were dried in the speed vac for 2 h to reach dryness, and then re-suspended in 0.2% formic acid.

#### Desalting

The re-suspended peptide samples were desalted with Zip Tips containing a C18 disk, concentrated and resuspended in aqueous 0.2% formic acid containing mass spectrometric “Hyper Reaction Monitoring” retention time peptide standards (iRT, Biognosys, Schlieren, Switzerland).

Briefly, samples were analyzed by reverse-phase HPLC-ESI-MS/MS using an Eksigent Ultra Plus nano-LC 2D HPLC system (Dublin, CA) with a cHiPLC system (Eksigent), which was directly connected to a quadrupole time-of-flight (QqTOF) TripleTOF 6600 mass spectrometer (SCIEX, Concord, CA). After injection, peptide mixtures were loaded onto a C18 pre-column chip (200 µm x 0.4 mm ChromXP C18-CL chip, 3 µm, 120 Å, SCIEX) and washed at 2 µl/min for 10 min with the loading solvent (H_2_O/0.1% formic acid) for desalting. Subsequently, peptides were transferred to the 75 µm x 15 cm ChromXP C18-CL chip, 3 µm, 120 Å, (SCIEX), and eluted at a flow rate of 300 nL/min with a 2-h gradient with aqueous and acetonitrile solvent buffers.

#### Acquisitions

Data-dependent acquisitions (for spectral library building): For peptide and protein identifications, the mass spectrometer was operated in data-dependent acquisition (DDA) mode, where the 30 most abundant precursor ions from the survey MS1 scan (250 msec) were isolated at 1 m/z resolution for collision-induced dissociation tandem mass spectrometry (CID-MS/MS, 100 msec per MS/MS, ‘high sensitivity’ product ion scan mode) using the Analyst 1.7 (build 96) software with a total cycle time of 3.3 sec as described (Christensen et al. 2018).

Data-independent acquisitions: For quantification, all peptide samples were analyzed by data-independent acquisition (DIA, e.g., SWATH), using 64 variable-width isolation windows (Collins et al. 2017; Schilling et al. 2017). The variable window width is adjusted according to the complexity of the typical MS1 ion current observed within a certain m/z range using a DIA ‘variable window method’ algorithm (more narrow windows were chosen in ‘busy’ m/z ranges, wide windows in m/z ranges with few eluting precursor ions). DIA acquisitions produce complex MS/MS spectra, which are a composite of all the analytes within each selected Q1 m/z window. The DIA cycle time of 3.2 sec included a 250 msec precursor ion scan, followed by 45 msec accumulation time for each of the 64 variable SWATH segments.

#### Mass-spectrometric data processing, quantification, and bioinformatics

Mass spectrometric DDAs were analyzed using the database search engine ProteinPilot (SCIEX 5.0, revision 4769) using the Paragon algorithm (5.0.0.0.4767) (Shilov et al. 2007), with search emphasis for ‘succinylation’. Using these database search engines results a MS/MS spectral library was generated in Skyline daily v20.2.1.404. The DIA/SWATH data were processed for relative quantification, comparing acylated peptide peak areas from various conditions. For the DIA/SWATH MS2 data sets, quantification was based on XICs of 6-10 MS/MS fragment ions, typically y- and b-ions, matching to specific peptides present in the spectral libraries. Significant changes were accepted at a 5% FDR (q-value < 0.05).

## Code and Data availability

Mass spectrometric raw data have been deposited to the MassIVE repository (MSV000089448) and are also available at ProteomeXchange (PXD033787). Code developed for this study is available at https://github.com/ccarrico/CoABindingSiteAnalyses.

## Acknowledgments

We acknowledge the NIDDK grant R24 DK085610 (Verdin), and we acknowledge the support of instrumentation from the NCRR shared instrumentation grant 1S10 OD016281 (Buck Institute). We thank Gary Howards and John Carroll for their help editing the manuscript and the figures, respectively.

## Figure Legends

**Supplementary Figure 1.**
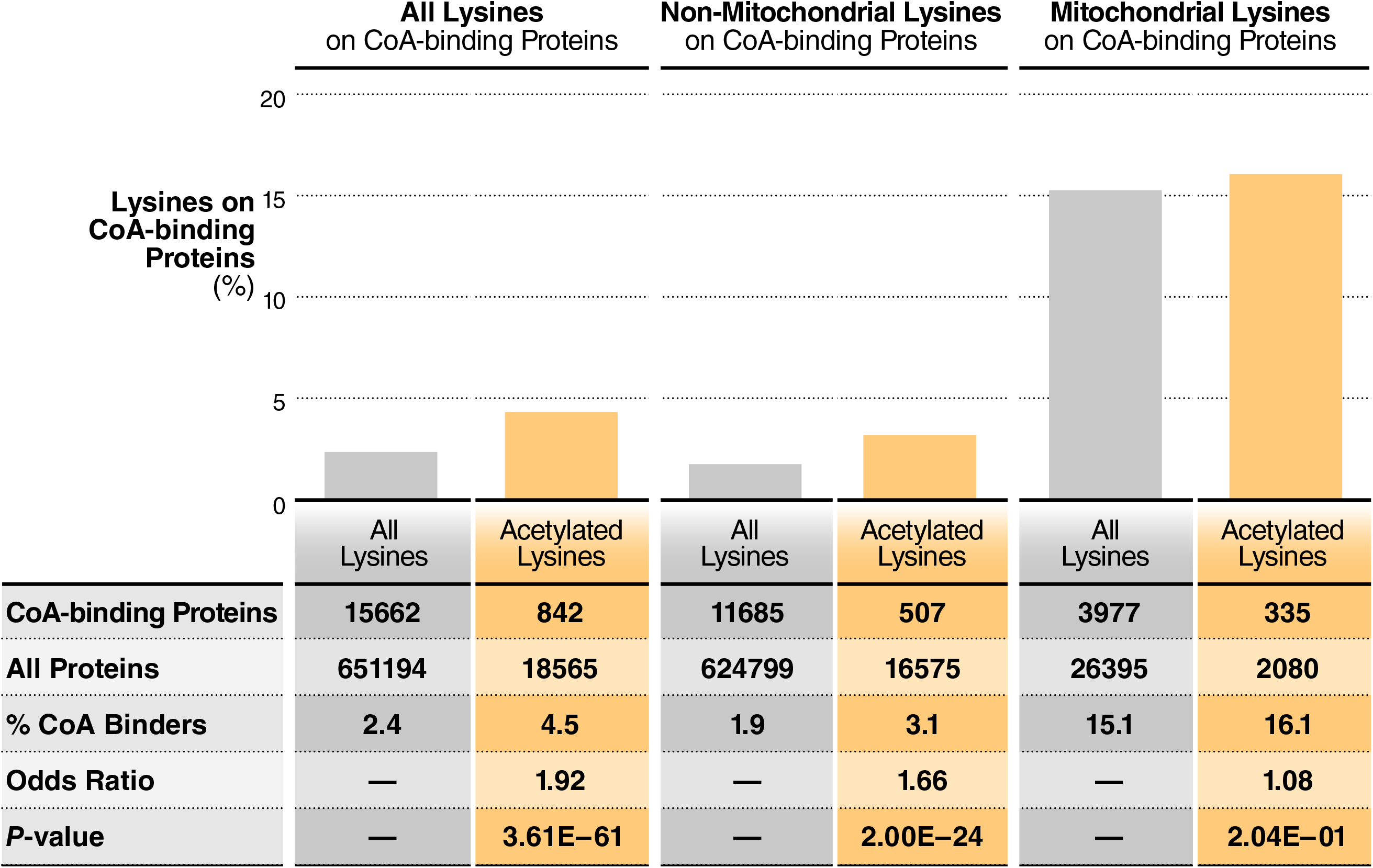
Enrichment of acetylated lysines on CoABPs analyzed in a whole-cell human acetylome dataset. Table showing the enrichment of acetylated lysines on CoABPs analyzed in a whole-cell acetylome dataset in human Hela cells. Statistical significances were calculated using Fisher’s exact test.

**Supplementary Table 1:** CoA-Binding Proteins in the Human and Mouse Proteomes

